# *Trichoderma reesei* complete genome sequence, repeat-induced point mutation and partitioning of CAZyme gene clusters

**DOI:** 10.1101/120071

**Authors:** Wan-Chen Li, Chien-Hao Huang, Chia-Ling Chen, Yu-Chien Chuang, Shu-Yun Tung, Ting-Fang Wang

## Abstract

*Trichoderma reesei* (Ascomycota, Pezizomycotina) QM6a is a model fungus for a broad spectrum of physiological phenomena, including plant cell wall degradation, industrial production of enzymes, light responses, conidiation, sexual development, polyketide biosynthesis and plant-fungal interactions. The genomes of QM6a and its high-enzyme producing mutants have been sequenced by second-generation-sequencing methods and are publicly available from the Joint Genome Institute (JGI). While these genome sequences have offered useful information for genomic and transcriptomic studies, their limitations and especially their short read lengths make them poorly suited for some particular biological problems, including assembly, genome-wide determination of chromosome architecture and genetic modification or engineering. We integrated Pacific Biosciences and Illumina sequencing platforms for the highest-quality genome assembly yet achieved, revealing seven telomere-to-telomere chromosomes (34,922,528 bp; 10877 genes) with 1630 newly-predicted genes and >1.5 Mb of new sequences. Most new sequences are located on AT-rich blocks, including 7 centromeres, 14 subtelomeres and 2329 interspersed AT-rich blocks. The seven QM6a centromeres separately consist of 24 conserved repeats and 37 putative centromere-encoded genes. These findings open up a new perspective for future centromere and chromosome architecture studies. Next, we demonstrate that sexual crossing readily induced cytosine-to-thymine point mutations on both tandem and unlinked duplicated sequences. We also show by bioinformatic analysis that *Trichoderma reesei* has evolved a robust repeat-induced point mutation (RIP) system to accumulate AT-rich sequences, with longer AT-rich blocks having more RIP mutations. The widespread distribution of AT-rich blocks correlates genome-wide partitions with gene clusters, explaining why clustering of genes has been reported to not influence gene expression in *Trichoderma reesei*. Compartmentation of ancestral gene clusters by AT-rich blocks might promote flexibilities that are evolutionarily advantageous in this fungus’ soil habitats and other natural environments. Our analyses, together with the complete genome sequence, provide a better blueprint for biotechnological and industrial applications.

## Background

*Trichoderma* is a fungal genus in soils and many other natural environments. *Trichoderma reesei* (syn. *Hypocrea jecorina*) is a widely used model organism for plant cell wall degradation and industrial enzyme production. The natural strain QM6a (ATCC13631) was first isolated from the Solomon Islands during the Second World War [1]. High enzyme producers (e.g., QM9414 and RUT-C30) were artificially generated from QM6a for industrial applications over the last 70 years [2-6].

*Trichoderma reesei* undergoes a heterothallic reproductive cycle and generates fruiting bodies (stromata) containing linear asci with 16 ascospores [7]. These 16 ascospores are generated via meiosis and two rounds of postmeiotic mitosis [8]. When placed under favorable conditions, ascospores germinate to form vegetative mycelia and produce asexual spores (i.e., conidia). Sexual development of the *Trichoderma reesei* CBS999.97 wild-isolate strain produced two haploid strains, CBS999.97(*MAT1-1*, F/X) and CBS999.97(*MAT1-2*, M/33). The two ancestral scaffolds (M and 33) in CBS999.97(*MAT1-2*, M/33) underwent an unequal translocation to form two new scaffolds (F and X) in CBS999.97(*MAT1-1*, F/X) [8]. Like CBS999.97(*MAT1-2*, M/33), QM6a has a *MAT1-2* mating type locus and two ancestral scaffolds M and 33. Due to chromosome heterozygosity and meiotic recombination, sexual crossing of CBS999.97(*MAT1-1*, F/X) with CBS999.97(*MAT1-2*, M/33) or QM6a often (>90%) generates segmentally aneuploid (SAN) progeny [8]. The CBS999.97 wild-isolate strain is also an excellent fungal model for light responses since light variation greatly affects its sexual development and conidiation [7, 9, 10]. Constant light promotes conidiation and completely inhibits stromata formation, whereas total darkness causes a slowdown of the growth of stromata [10].

The genomes of several *Trichoderma* species have been sequenced and are publicly available from the Joint Genome Institute of the US Department of Energy. The QM6a-v2.0 draft genome (33.4 Mb) contains 87 scaffolds and 9129 predicted genes [11]. The RUT-C30-v1.0 draft genome (32.7 Mb) has 182 scaffolds and 9852 predicted genes [12]. The genomes of *Trichoderma atroviride* and *Trichoderma virens* were thought to be larger than QM6a-v2.0, with sizes of 36.1 and 38.8 Mb, respectively, versus 34.1 MB for QM6a, both encoding more than 2,000 additional predicted genes [4, 13]. These genomes have been used to identify key genes involved in some important biological processes [4, 13-15], e.g., the transcriptional factors and transporters that control induction and expression of carbohydrate-active enzymes (CAZymes) and the plant cell wall degradation enzymes. Using QM6a as a reference, it has also been reported that QM9414 and RUT-C30 might carry multiple alternations, including rearrangements, point mutations, insertions and deletions [12, 16]. Recently, a group of *Trichoderma* researchers have collectively annotated and compared ∼30% genes in the JGI genomes of *Trichoderma reesei, Trichoderma atroviride* and *Trichoderma virens* [4].

To reveal gene order and dynamic gene expression at the chromosome level, a genome-wide chromosome conformation capture method (referred herein as “HiC”) had been applied to close the gaps between the QM6a-v2.0 scaffolds. The HiC draft genome revealed seven super-scaffolds and four short contigs [17]. Druzhinina et al. then applied the QM6a-HiC draft to annotate 9151 (not 9194) predicted genes. A third of the putative CAZyme genes occurred in loose clusters that also contained a high number of genes encoding small secreted cysteine-rich proteins (SSCPs). Five CAZyme gene clusters are located close to chromosomal ends. These subtelomeric areas are also enriched in genes involved in conidiation, iron scavenging and interactions with other fungi, such as secreted protease genes, amino acid transporter genes, gene clusters for polyketide synthases (PKS), non-ribosomal peptide synthase (NRPS) and PKS-NRPS fusion proteins [18]. The QM6a-HiC annotation (http://trichocode.com/index.php/t-reesei) [18] became publicly available in January, 2017. Because it does not provide any expectation values (E) for the BLAST sequence alignments, its reliability needs to be further confirmed.

Strictly speaking, the HiC draft genome is far from equivalent to a complete genome sequence. We noticed that both the QM6a-v2.0 and QM6a-HiC drafts lack several evolutionarily-conserved genes that are ubiquitiously expressed in almost all studied eukaryotic organisms, including the recA family protein Rad51 and the DNA repair protein Rad50. Therefore, we applied both second and third generation sequencing (SGS and TGS) platforms to resequence the QM6a genome. The Single Molecule Real Time (SMRT) sequencing method developed by Pacific BioSciences (PacBio) offers much longer reads of up to 60 kb [19]. After error correction with short and high-quality Illumina-MiSeq sequencing reads, the PacBio long reads were assembled into seven telomere-to-telomere chromosomes. Our high-quality genome sequence provides a large quantity of new information to facilitate functional and comparative studies of this industrially important workhorse fungus.

## Results

### Resequencing the QM6a genome

The QM6a genome was resequenced using seven SMRT cells on the PacBio RSII platform. Following extraction of reads with *Trichoderma reesei*-only sequences, we recovered the longest raw reads (≥16□kb) with up to 80x coverage, totaling 3,397,762,180 bp. The hierarchical genome assembly process program [20] was used to generate a preliminary PacBio draft with seven superscaffolds, a short unitig and 1.8 kb of contaminating DNA (Additional File 1: Table A1). This short unitig was completely identical in nucleotide sequence to the QM9414 mitochondrial genome (42,139bp; NC_003388.1) [21], indicating that the error rate for the preliminary PacBio draft was extremely low (<0.0024%).

For error correction, the Ilumina MiSeq 300 bp paired-end reads (6.8 Gb) were collected and trimmed. Reads (67.85%) with a quality score threshold (Q) greater than 30 were retained (Additional File 1: Table A2 and A3). The final assembly data contains a circular mitochondrial genome (42,139 bp) and seven unitigs (34,922,528 bp) (Additional File 1: Table A3). We highlight that there were no sequence ambiguities or unidentified bases (Ns) (Table 1). In contrast, the QM6a-v2.0 draft genome was 33,453,791 bp and had 48,252 Ns, whereas the HiC draft was 33,395,328 bp and had 42,879 Ns.

The seven superscaffolds closely match (if not being completely identical to) the full-length chromosomes because all of their termini capture typical telomeric sequences (i.e., TTAGGG at 3’-termini and the reverse complement CCCTAA at 5’-termini) [11] with up to 14 repeats (Additional File 1: Table A3). We categorized these telomere-to-telomere chromosomes with Roman numerals (ChI-ChVII), from largest to smallest. Genetically-defined linkage groups were designated alphabetically (A-G), and linkage group arms are designated L for the short (left) arm at 5’ termini and R for the long (right) arm at 3’ termini (Figure 1). The complete QM6a genome sequences have been submitted to NCBI (accession number CP016232-CP016238).

**Figure 1.**
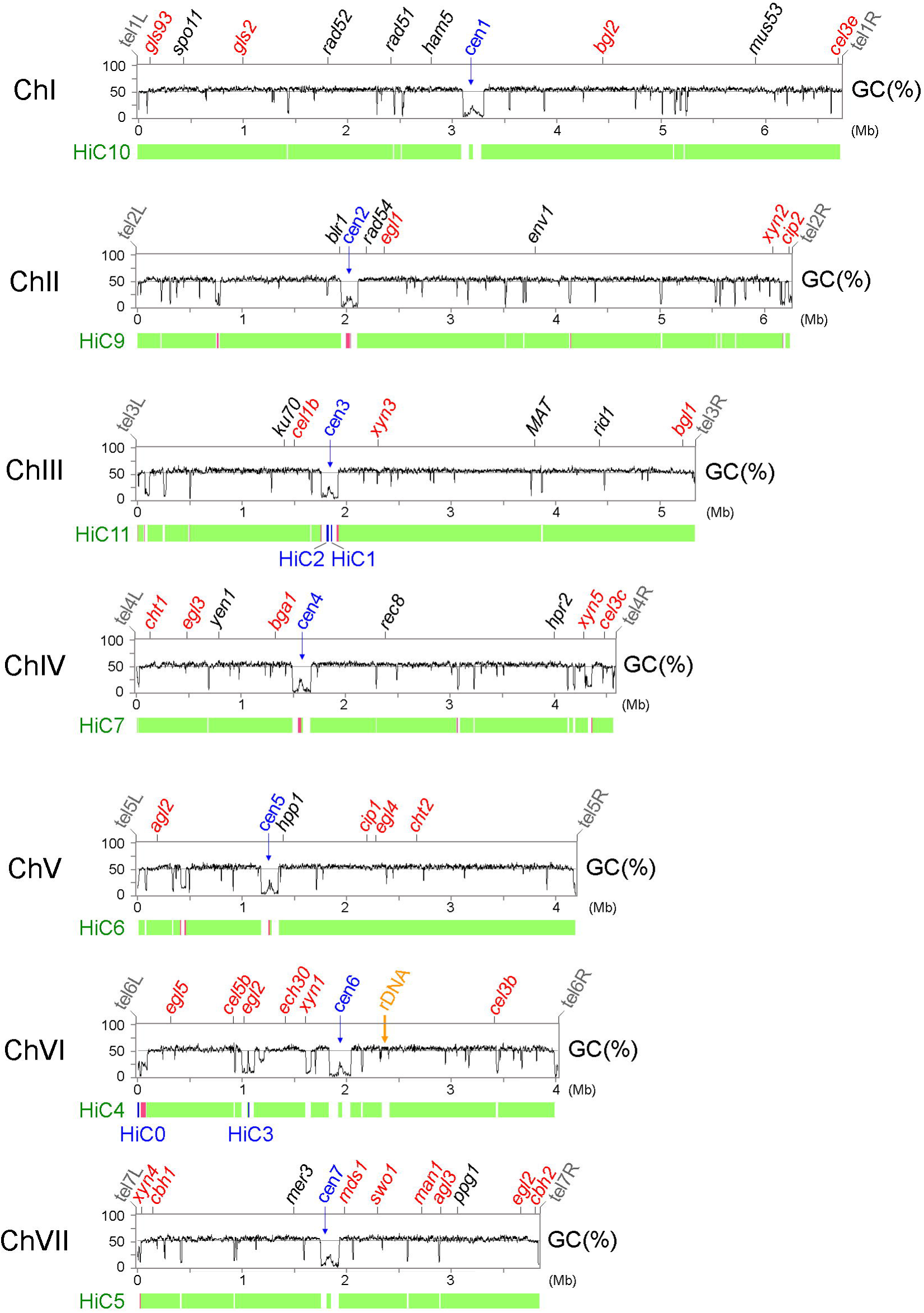
The complete QM6a genome compared to the HiC draft genome. The top tracks represent the graphs of GC contents (window size 5000 bp) of seven telomere-to-telomere chromosomes (ChI-ChVII). The seven centromeres (*cen1-cen7;* in blue) are located at the longest AT-rich blocks in each chromosome. The telomeres (*tel*) of the right (*R*) and left (*L*) arms in each chromosome are indicated (in grey). The CAZyme genes (in red) and several genes involved in DNA repair, light response and sexual development (in black) are indicated. The rDNA (18S-5.8S-26S) locus on ChVI is indicated in orange. The bottom tracks represent the seven superscaffolds (in green) and four short contigs (in blue) of the HiC draft genome. Chromosomal regions with incorrect orientation (i.e., inversion errors) are indicated in pink.

### The complete QM6a genome compared to two earlier draft genomes

By mapping the MiSeq reads to the QM6a-v2.0 draft, we found that ten scaffolds in the QM6a-v2.0 draft were not genuine *Trichoderma reesei* sequences (Additional File 1: Tables A5). For this reason, 16 previously annotated genes (Additional File 1: Tables A6) in QM6a-v2.0 and QM6a-HiC might not be authentic QM6a genes. In addition, there were numerous sequencing and assembly errors in QM6a-v2.0. The most prominent assembly error was the first and longest scaffold, wherein the 5’ portion (∼2.46 Mb) was mapped to ChV and the 3’ portion (∼1.79 Mb) to ChIV (Additional File 1: Table A7).

Our complete QM6a genome sequence also covers all seven superscaffolds and the four short contigs of QM6a-HiC [17]. QM6a-HiC contains a large quantity of sequence and/or assembly errors, particularly those sequences (e.g., telomeric repeats) close to both termini of the seven superscaffolds. The four short HiC contigs are all located at chromosome regions with low guanine(G)-cytosine(C) contents. HiC0 is located close to *tel6L* (the left telomere of ChVI), HiC1 and HiC2 at *cen3* (the centromere of ChIII), and HiC3 in an interspersed AT-rich block on the right arm of ChVI. There were at least 18 inversion errors in the HiC draft genome (Figure 1 and Additional File 1: Table A8). Eight of these inversion errors could account for the failure to connect these four short contigs to the corresponding superscaffolds during the HiC experiments [17].

The HiC experiments also resulted in incorrect assembly at the rDNA locus. Using Southern hybridization, we confirmed that the right arm of ChVI harbors the large rDNA locus with nine tandem “head-to-tail” repeats. Each repeat contains an 18S–5.8S–26S rRNA gene cluster and a non-transcribed intergenic spacer (IGS) (Figure 2 and Additional File 1: Table A9). This result approaches the theoretical limit for mapping results using Illumina Miseq short reads, i.e., 200-260x coverage at the rDNA locus versus 25-30x coverage along the entire chromosome (Figure 3). It is worth noting that there are 175-200 copies of the large rDNA tandem repeats in *Neurospora crassa* [22].

**Figure 2.**
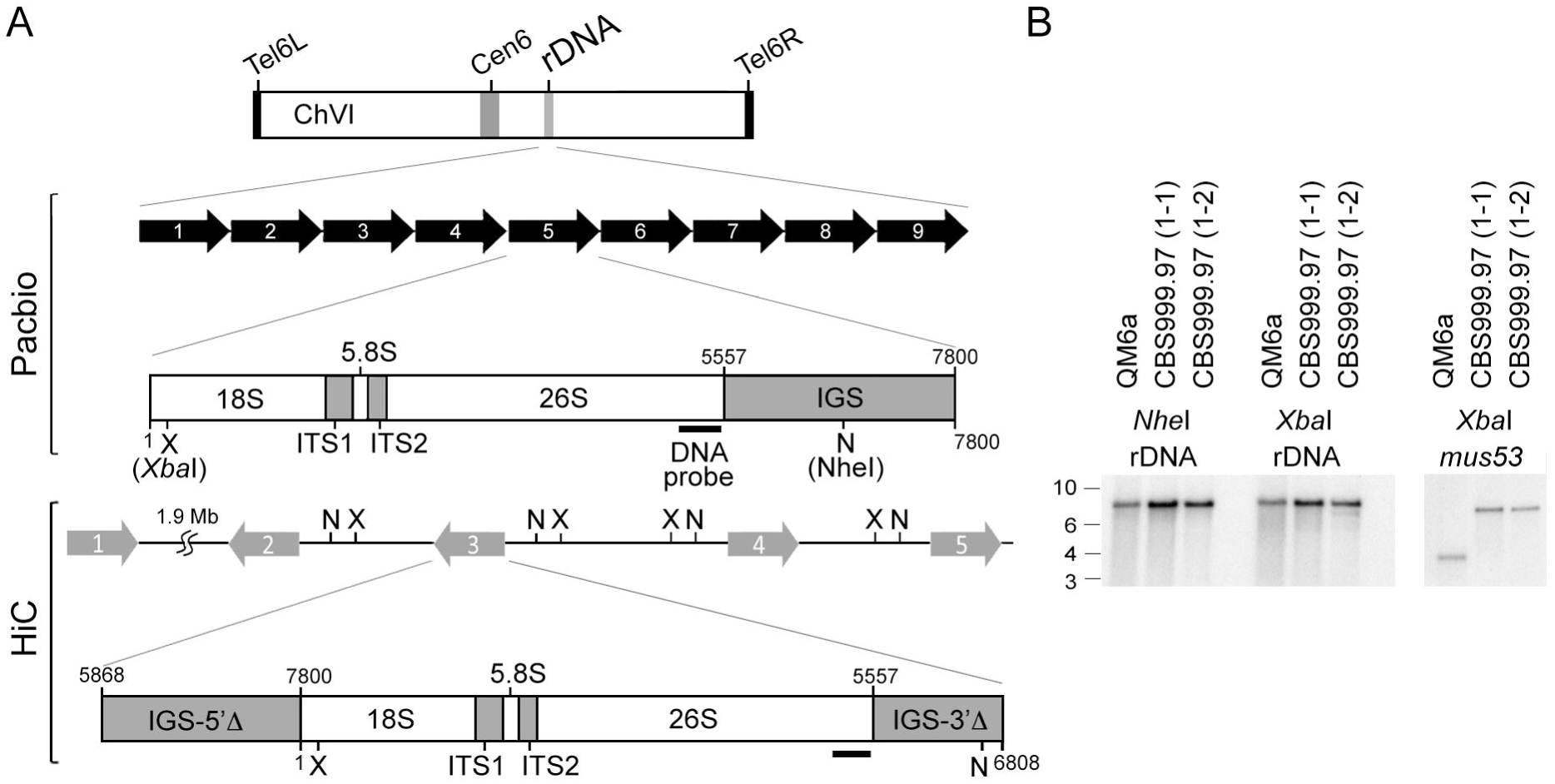
The rDNA locus. (**A**) Organization of the rDNA locus. The top panel illustrates the nine tandem “head-to-tail” repeats revealed by the PacBio RSII platform. Each repeat contains an 18S–5.8S–26S rRNA gene cluster and a full-length non-transcribed intergenic spacer (IGS). The bottom panel shows the five repeats in the HiC draft genome. Each repeat has an 18S-5.8S-26S rRNA gene cluster and two truncated IGSs (IGS-5’Δ and IGS-3’Δ). The location of restriction enzymes *Xbal* (X) and *Nhel* (N) are indicted. (**B**) Southern hybridization. Genomic DNA (1 μg) was isolated from three different wild isolate strains: QM6a, CBS999.97(1-1) and CBS999.97(1-2). After digestion with *XbaI* (X) or *NheI* (N), the genomic DNA was subjected to agarose gel electrophoresis, Southern blotting and hybridization with a 28S rDNA probe (**A**) or a *mus53* probe (as the DNA loading control). The *mus53* gene encodes the DNA ligase IV protein and there is only one copy of the *mus53* gene in the *Trichoderma reesei* genome [78].

**Figure 3.**
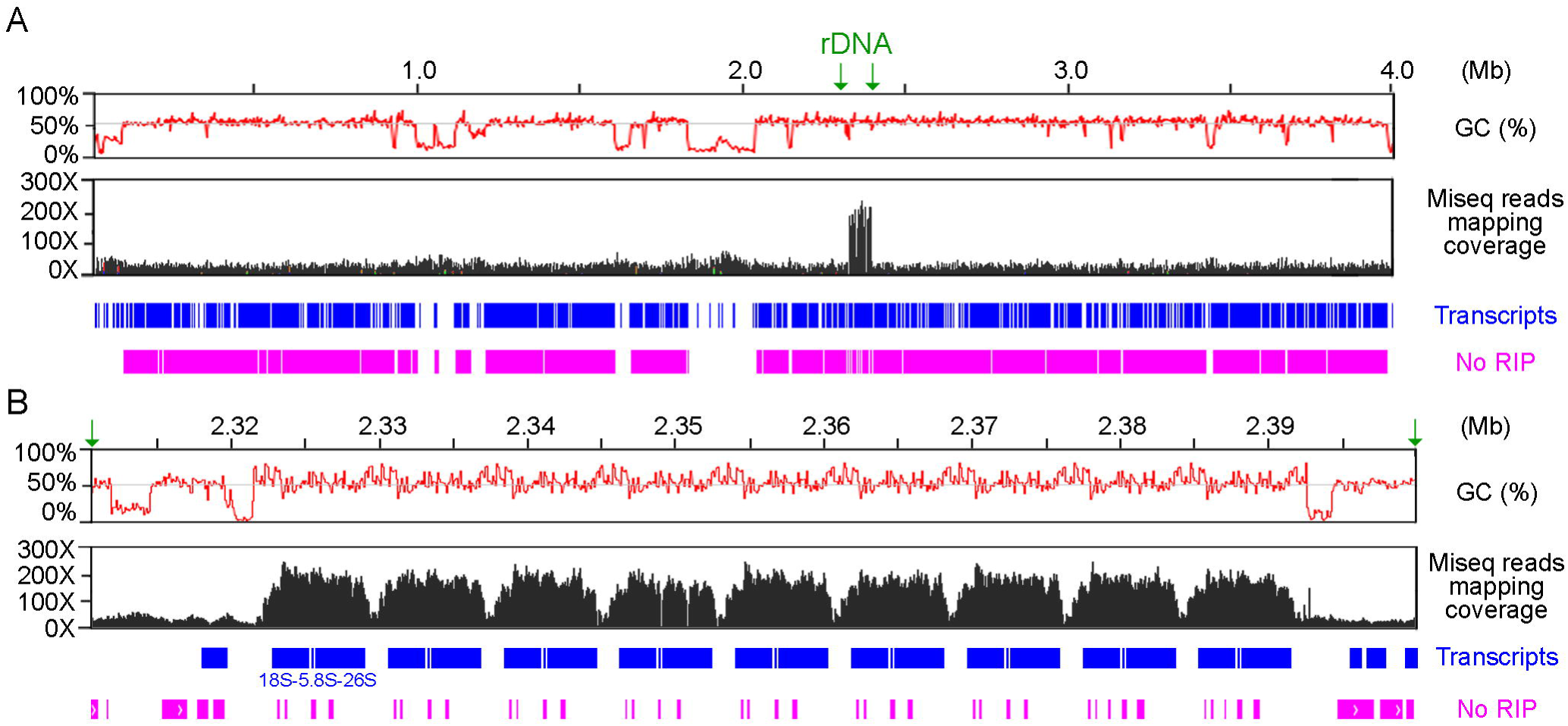
The repetitive features of a representative QM6a chromosome (ChIV). The top tracks represent the graphs of GC contents (in red; window size 100 bp), the mapping coverage of Illumina-MiSeq reads (in black), predicted genes (in blue) and the sequences not affected by RIP (No RIP; in pink) along the entire ChIV (**A**) and the rDNA locus (**B**). The mapping coverage of Illumina-MiSeq reads for overall genomic DNA is 25-30X, whereas that specifically for the rDNA locus is 200-260X. The RICAL program was used to predict the sequences mutated by RIP using default settings, and the sequences not affected by RIP are shown (No RIP; in pink).

Thus, the complete QM6a genome sequence has uncovered many sequencing and assembly errors in QM6a-v2.0 and QM6a-HiC. We suggest that caution should be exercised in applying HiC or the chromosome conformation capture method for gapclosing draft genome sequences produced by SGS technology and for other genomic analyses involved in AT-rich and repetitive sequences.

### QM6a compared to high enzyme producers

The high-quality QM6a genome sequence we provide here is a better scaffold to order and orient contigs of other *Trichoderma* draft genomes previously generated by SGS technologies. It has been reported that there might be five or eleven potential translocations in RUT-C30 [12, 16]. A BLAST search revealed that there are only three promising translocations in RUT-C30: ChV to ChII (the first scaffold of RUT-C30), ChIII to ChI (the second scaffold of RUT-C30), and ChIII to ChIV (the fifth scaffold of RUT-C30). Theoretically, these three translocations in RUT-C30, together with the one translocation we identified in CBS999.97(*MAT1-1*, F/X), are sufficient to account for the large quantity of inviable SAN ascospores generated from sexually crossing RUT-C30 with CBS999.97(*MAT1-1*, F/X) [8, 23]. There are also three short ectopic insertions in RUT-C30 including the second scaffold (34 bp), the fifth scaffold (231 bp) and the ninth scaffold (2,655 bp) (Additional File 1: Table A10). These three ectopic insertions are more likely due to sequence duplication or assembly errors.

### Genome reannotation

We applied four different approaches to genome reannotation (Materials and Methods), including the use of all (28748) *Trichoderma reesei* proteins from National Center for Biotechnology Information (NCBI), QM6a-v2.0, RUT-C30-v1.0 and two publicly-available transcriptome datasets [24, 25]. We annotated 1630 newly-predicted QM6a genes, including 70 tRNA genes and 23 5S-rDNA genes (Additional File 2: Tables B1 and B2). The average length of all 10876 QM6a genes is 1579 bp. Their average GC content (56.5%) is ∼5.5% higher than that of the entire QM6a genome. Among 1515 new protein-encoding genes, 679 have been annotated in RUT-C30-v1.0. Most of them encode novel or hypothetical proteins, and only 120 and 285 newly-predicted genes encode protein products that have homologs in *Saccharomyces cerevisiae* and *Neurospora crassa*, respectively. It is worth noting that we annotated several essential or biologically important genes, including six DNA repair genes (*rad50, sae2/com1, rad51, rad57, srs2, rrm3* and *pif1*), an essential component of sister chromatin cohesion complex (*smc1*), a key autophagy gene (*atg11*), two cell division cycle genes (*cdc4* and *cdc15*) and two mitochondrial genes (*sod2* and *tom7*) (Additional File 1: Table A11 and Additional File 2: Tables B2 and B3). These evolutionarily conserved genes had never been annotated in QM6a-v2.0 or QM6a-HiC [4, 11,18]. The complete genome sequence and set of genes we provide here can serve as a better guide for further experiments, especially for global approaches to evolution, biological functions and industrial applications.

We categorized all QM6a genes with a non-italicized uppercase letter, a number and a letter: Tr (for *Trichoderma reesei*); A, B to G (for chromosome I, II through VII); a number corresponding to the order of the transcripts (counting from the left telomere); and W or C to designate the Watson or Crick strand (the Watson strand is 5’→ 3’ left telomere to right telomere); for example, the 100th gene from the left telomere of chromosome I is TrA0100C (Additional File 2).

### Comparative transcriptome analysis

Next, we applied TopHat—a bioinformatic sequence analysis package tool—to map and count the SGS reads of all annotated genes and then to determine the values of Transcripts Per kilobase Million (TPM) [26]. Compared to RPKM (Reads Per Kilobase Million) and FPKM (Fragments Per Kilobase Million), TPM is a stable and reliable RNA-seq expression unit across experiments [27]. All reads mapped to the rRNA and tRNA genes were excluded before calculating TPMs (Additional File 2: Tables B1).

Our results confirmed those reported by [24] that 35 CAZyme genes and 27 non-CAZyme genes were highly induced (≥ 20-fold) by straw substrate in QM6a (Additional File 3: Table C1). In addition, straw also upregulated (≥ 20-fold) 210 previously annotated genes in QM6a-v2.0, including *xyr1* (xylanase regulator 1; ≥ 100-fold), 38 CAZyme genes, 1 NRPS gene, the mating type gene *mat1-2-1* [7] and hybrid-type peptide pheromone precursor 1 (*hpp1*) [28] (Additional File 3: Table C2). Many straw-induced genes in QM6a [24] were shown to be differentially regulated in response to cellulose and sophorose in QM9414 [25] (Additional File 3: Table C1 and C2).

Among the 1535 new protein-encoding genes, there are 39 straw-induced (≥ 20-fold) genes in QM6a and 2 cellulose-induced genes (≥ 20-fold) in QM9414 (Additional File 3: Table C3). One hundred and forty four new QM6a genes are also significantly upregulated (5-20 fold) by straw (Additional File 3: Table C4). For example, TrC1345C, a new straw-induced gene, encodes the homolog of *S. cerevisiae* Cat8 zinc-cluster transcription factor. Cat8 is involved in gluconeogenesis, the glyoxylate cycle and ethanol utilization, and it is necessary for derepression of a variety of genes under non-fermentative growth conditions (e.g., diauxic shift and sporulation) [29]. Further research on this transcriptional factor might help to enhance cellulolytic enzyme production. We also identified nine new genes with TPM values in QM6a that are much higher than those in QM9414 (Additional File 3: Table C5). These genes are likely mutated or even deleted in QM9414.

### Wide distribution of AT-rich blocks along seven QM6a chromosomes

To establish the link between DNA sequence and chromatin architecture, the local GC contents along each chromosome were calculated using a 0.5 kb sliding window. We identified 2349 AT-rich chromosome blocks with GC contents ≥12% and ≥6% lower than the average GC content of the all predicted genes (56.5%) and the entire QM6a genome (51.1%), respectively (Table 2).

The most prominent or longest AT-rich blocks in each chromosome are the centromeres, ranging from 162.5 kb (*cen2*) to 208.5 kb (*cen6*) (Figure 4 and Additional File 3: Table A2). The longest AT-rich blocks in each *Neurospora crassa* chromosome are also centromeres [30]. A BLAST search revealed that the seven QM6a centromeres collectively harbored 24 conserved sequences (≥90% identity; maximum length 8625 bp and minimum length 4847 bp) with a copy number per chromosome ranging from one to five (Additional File 4). These conserved sequences are centromere-specific and highly AT-rich, perhaps representing centromeric repeats.

**Figure 4.**
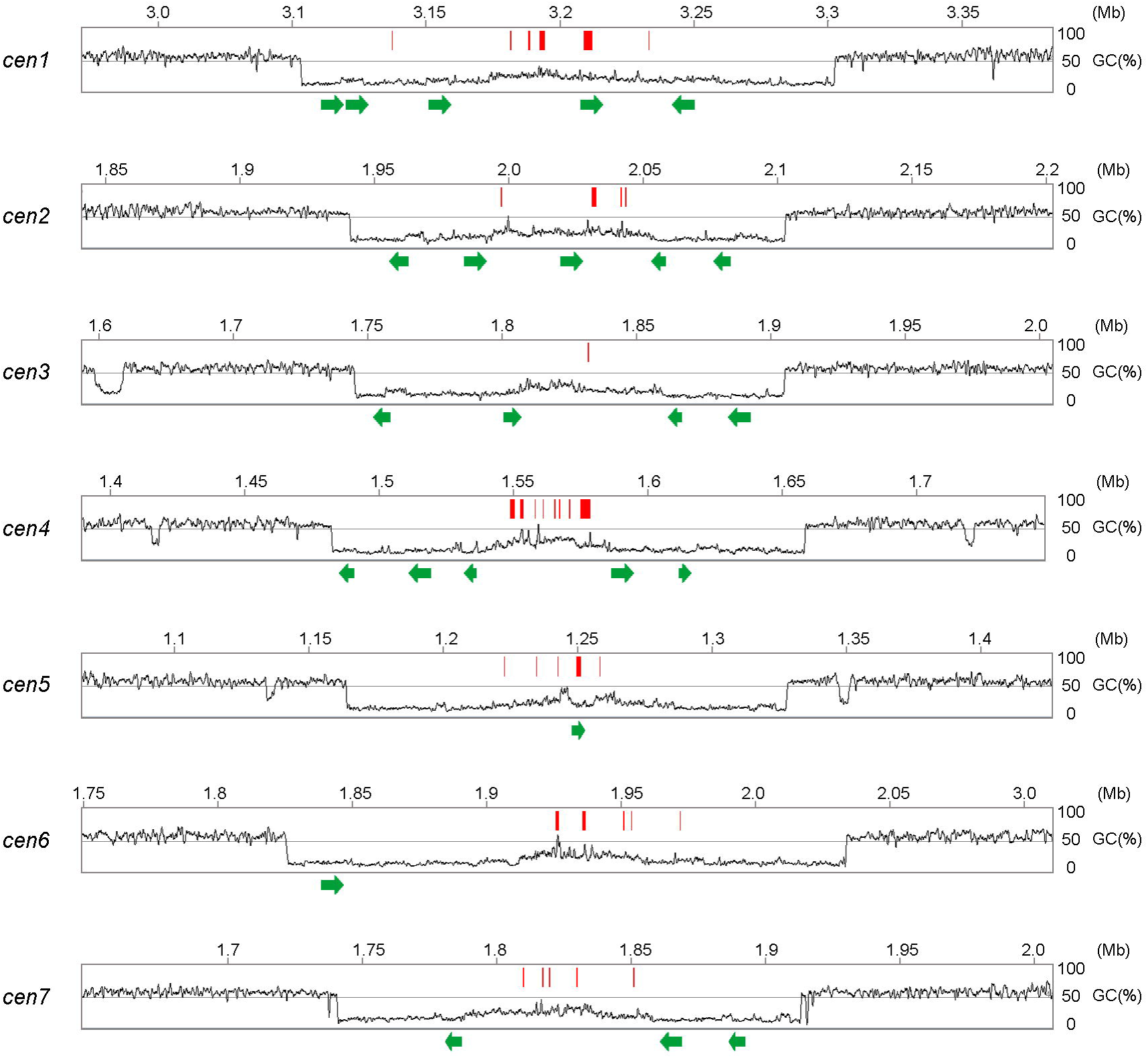
QM6a centromeres. The tracks represent the graphs of GC contents of seven centromeres (*cen1-cen7*) and the corresponding pericentromeric regions. The centromere-encoded genes are indicated by red bars and the conserved centromeric repeats are indicated by green arrows.

The seven QM6a centromeres also separately encode 13 previously-annotated genes and 24 newly-predicted genes (Additional File 3: Table C6). Centromere-encoded transcripts are known to be integral components of the genomes of mammals, higher plants and the fission yeast *Schizosaccharomyces pombe* [31-34]. Copy numbers of these 37 centromere-encoded transcripts were relatively low; their expression (TPM > 0) could only be detected in QM9414, but not in QM6a (Additional File 3: Table C6). The QM9414 transcriptomic data from the Illumina HiSeq 2000 platform [25] apparently had better sequencing depth than the QM6a transcriptomic data from the SOLiD platform [24]. It will be of importance to investigate whether these 24 centromeric repeats and 37 centromere-encoded genes are involved in centromere integrity and chromosome segregation fidelity in *Trichoderma reesei*.

The AT-rich chromosomal blocks next to the 14 telomeres are subtelomeres, with the shortest being ∼1 kb and the longest up to 87 kb (Additional File 1: Table A3). Other than centromeres and subtelomeres, there are 2328 interspersed AT-rich blocks (Table 2). On average, there are only five genes between two neighboring interspersed AT-rich blocks. The biological relevance of these interspersed AT-rich blocks remains to be elucidated (see below). It is worth noting that both the 5’ and 3’ flanking sequences of the rDNA locus contain a 2 kb AT-rich block (Figure 3). We postulate that these two AT-rich blocks might be involved in regulating nucleolar organization, rRNA transcription, rDNA copy homeostasis and prevention of repeat-induced point mutation (RIP) (see below).

### Comparative genomic analyses of chromosome architectures

Next, we compared the QM6a genome with 11 publicly-available and well-assembled fungal genomes for their AT-rich blocks. We highlight that all these fungal genomes contain ≤ 0.11% unresolved or unknown bases (Ns) (Additional File 1: Table A12). Unresolved bases jeopardize the integrity of *in silico* genomic analysis.

All of these 12 fungal genomes have many short AT-rich blocks with lengths of 0.5-3 kb (Table 2). In *Saccharomyces cerevisiae*, these short AT-rich blocks correlate with the convergent intergenic regions and the pericentromeres that associate with cohesin (an evolutionarily-conserved protein complex that functions to hold a pair of sister chromatids during mitosis and meiosis). The average distance between neighboring cohesin-binding sites along yeast chromosome arms is 10-15 kb, which is compatible with the observed localized oscillations in base composition [35-37]. It has been suggested that sister chromatid connections via cohesin complexes occur preferentially at the chromosome axes, the bases of intrachromosomal loops or topologically-associated domains (TADs) [35, 38-40]. Expression of genes within TADs is somewhat correlated [41-45]. Intriguingly, the average distance between neighboring AT-rich blocks along QM6a chromosomal arms is ∼13 kb. We suggest that these AT-rich blocks might be functionally associated with chromosomal loading of cohesin in *Trichoderma reesei*.

We were able to categorize these 12 fungal genomes into three different groups according to: (1) the average AT content, (2) the genome-wide distribution of local AT content, and (3) the number of long (≥ 3kb) AT-rich blocks. For example, in the QM6a genome, there were 167 AT-rich blocks with length ≥ 3 kb (Table 2 and Figure 6).

The Group I fungi consist of six filamentous ascomycetes (Pezizomycotina), including QM6a, *Neurospora crassa* (OR74A), *Penicillium chrysogenum* (P2niaD18), *Mycosphaerella graminicola* (IPO323), *Fusarium fujikuroi* (IMI 58289), and *Aspergillus nidulans* (FGSC A4). Their genomes not only have similar average AT contents (∼50%) but also display biphasic local AT content distributions due to the presence of many longer (≥ 3kb) AT-rich blocks (Table 2 and Figure 6A, top two panels). In *Neurospora crassa*, AT-rich blocks are DIM-3 (importin α)-dependent constitutive heterochromatins with transposon relicts and trimethylated histone 3 at lysine 9 (H3K9me3). Genome organization in *Neurospora crassa* nuclei is largely defined by constitutive heterochromatins via strong intra- and inter-chromosomal contacts [46, 47].

Group II are three basidiomycetes, i.e., *Ustilago maydis* (521), *Coprinopsis cinerea* (Okayama 7#130) and *Cryptococcus neoformans* (JEC21). Their average AT contents are ∼50%, similar to those of Group I fungal genomes (Table 2). The Group II fungal genomes display relatively normal local AT content distribution (Figure 5A, the lowest panel) due to the absence of longer AT-rich blocks (Figure 5B).

**Figure 5.**
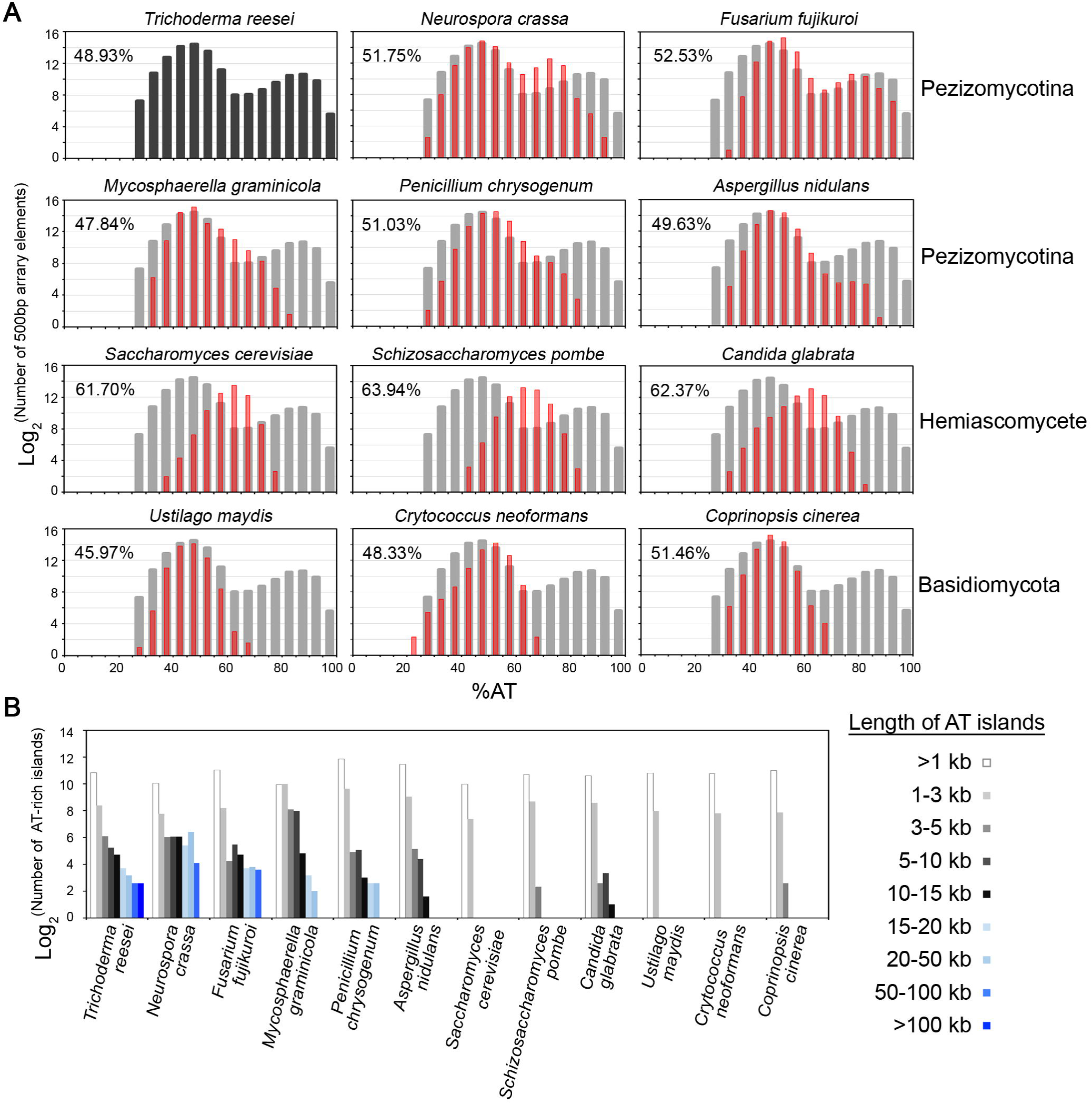
Comparative analysis of chromosome architecture for 12 different fungal genomes. (**A**) The AT content for each array element (500 bp) was calculated and put into bins of 5% intervals (black or gray bars for QM6a and red for other fungi, y-axis). The average AT content of each fungal genome is shown. (**B**) Numbers of AT-rich blocks of different lengths are shown. We were able to categorize these 12 fungal genomes into three different groups according to: (1) their average AT contents, (2) the genome-wide distribution of local AT content, and (3) the number of long (≥ 3kb) AT-rich blocks.

Group III are three hemiascomycete yeasts, *Saccharomyces cerevisiae* (S288C), *Schizosaccharomyces pombe* (972h-) and *Candida glabrata* (CBS138). Their average AT contents (62-64%) are 12-14% higher than those of Group I and Group II fungi (Table 2). These three yeast genomes also display a relatively normal distribution of AT local content (Figure 5A, the lowest panel) and have no or very few longer AT-rich blocks (Figure 5B).

### Repeat-Induced Point mutation (RIP) is identified in *Trichoderma reesei*

The high number of long AT-rich blocks in the Group I fungal genomes might be correlated with RIP: a phenomena originally discovered in *Neurospora crassa* at a premeiotic stage during sexual development [46]. The process of RIP requires a specialized cytosine methyltransferase gene *rid-1* (RIP-defective) and induces cytosine-to-thymine (C-to-T) point mutations in a homology-dependent manner [48, 49]. Several previous studies found no evidence of RIP in any of the Group II or Group III fungi we investigated here [50-52]. RIP has been documented in *Penicillium chrysogenum* [53], *Fusarium fujikuroi* [54], *Mycosphaerella graminicola* [55] and *Aspergillus nidulans* [56], but it has never been experimentally demonstrated in any *Trichoderma* species [18, 57]. Intriguingly, the QM6a complete genome encodes almost all proteins known to be involved in RIP and DNA methylation in *Neurospora crassa*, including *rid1* (TrC1298W) [57], *dim2* (TrB0908W; DNA methyltransferase), *dim3* (TrE0517W; importin α), *dim5* (TrB0159C; H3K9 methyltransferase) [13], *dim7* (TrD0784W), *dim8* (TrG0915W), *dim9* (Trc0267C) and *hpo* (TrB1406W; HP1) (Additional File 2: Tables B1 and B2).

We carried out both bioinformatic and molecular genetic analyses to determine whether RIP could operate in *Trichoderma reesei*. The RIPCAL software tool was applied to compare differences in the extent of RIP mutations of different sequences by determining two widely-used RIP indices: TpA/ApT and [(CpA + TpG) / (ApC + GpT)] [58, 59]. Higher values of TpA/ApT and lower values of [(CpA + TpG) / (ApC + GpT)] indicate stronger RIP responses [60, 61]. We found that the hierarchy for RIP in QM6a is the mating type gene *MAT1-2-1* < all predicted genes < the whole QM6a genome < the large rDNA tandem repeats (18S-5.8S-26S) < 5S rDNAs. It has been reported that the large rDNA tandem repeats and the 5S rDNAs in *Neurospora crassa* survived RIPs due to either nucleolar sequestration [62] or their smaller size [60], respectively. The RIP indices of the mating gene (*MAT1-2-1 or MFa*), all predicted genes and the whole genome suggest that the RIP respones in QM6a are as robust as those in *Neurospora crassa* (Additional File 1: Table A13). Thus, we suggest that *Trichoderma reesei* has evolved an RIP system similar to that of *Neurospora crassa*. Our results also reveal that the longer the AT-rich blocks the greater the extent of RIP mutations in QM6a (Additional File 1: Table A13), consistent with RIP mutations accumulating in AT-rich sequences.

Next, we carried out sexual crossing tests between four parental (F0) strains, including wild-type CBS999.97 [7], *blr1*Δ, *env1*Δ [9, 10] and *ku70*Δ [63], and then tested their F1 progeny for mutations in the full-length hygromycin-resistant (*hph*) genes present in the selection marker construct used for deletion of the corresponding *Trichoderma reesei* genes [10]. Progeny were isolated using the hexadecad dissection technique [8]. No *hph* sequence is present in the wild-type CBS999.97 F0 strains, whereas *env1*Δ and *ku70*Δ each carries one copy of full-length *hph*. In contrast, the *blr1*Δ mutant contains two tandem head-to-tail *hph* sequences resulting in repeats; one is a full-length *hph* and the other an N-terminal truncated *hph*-ΔN (Figure 6A). All of the *hph* and *hph*-ΔN alleles in the corresponding parental strains were confirmed first by genomic PCR and Sanger sequencing (see Additional File 5 for their nucleotide sequences) and then by Southern hybridization (Figure 6B).

**Figure 6.**
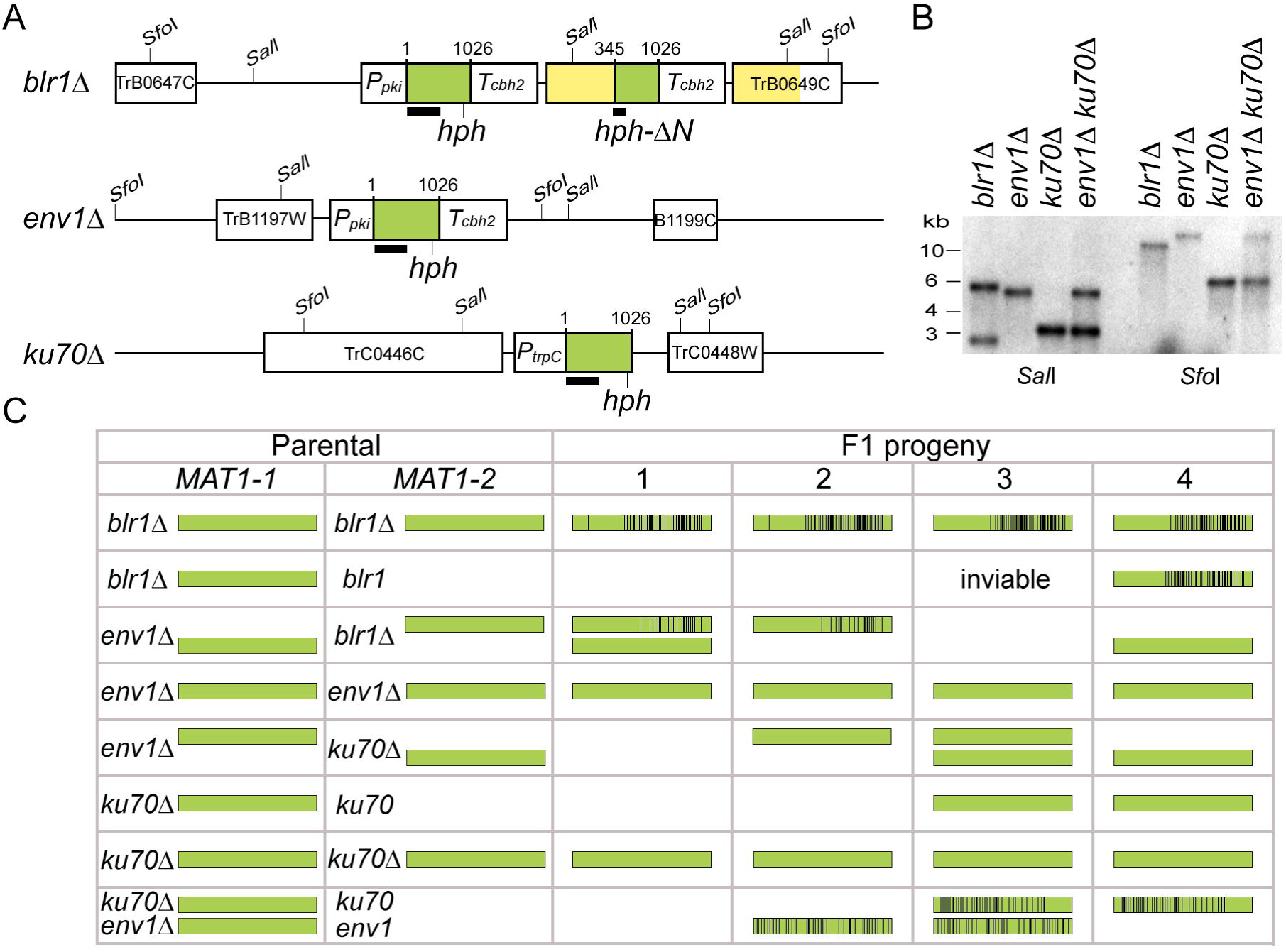
RIP in. *Trichoderma reesei*. (**A**) Schematic diagram of the gene deletion cassettes in the three mutants. Each cassette consists of three components; the dominant hygromycin B-resistant marker open reading frame (hph; green box), flanked by an upstream promoter (*P_pki_* or *P_trpC_*) and/or a downstream terminator (*T_cbh2_*). The full-length *hph* gene has 1026 bps. A truncated *hph-*Δ*N*_304-1026_*-T_cbh2_* cassette was spontaneously generated in *blr 1*Δ. during transformation. The dark line underneath *hph* represents the DNA probe used for Southern hybridization. The neighboring genes upstream and downstream of the deleted gene are indicated by white boxes and their protein identity numbers. In *blr*.Δ, there are two identical 12960 fragments (in yellow) and two copies of *T_cbh2_* (in white). (**B**) Southern hybridization. Genomic DNA of indicated strains was digested by *SalI* or *Sfo1*, and then subjected to Southern blot analysis using the *hph* DNA probe shown in (A). (**C**) Occurrence of RIP in the F1 progeny. The C-to-T point mutations in the respective full-length *hph* cassesttes (in green) are indicted by vertical black bars. Only the results of one representative sexual crossing experiment (n = 10) are shown. The full-length *hph* cassette in each progeny was amplified by polymerase chain reaction (PCR) and then sequenced by Sanger’s method.

After sexual crossing (n = 10), the F1 progeny displayed numerous C-to-T point mutations in all cases where two similar sequences were present in one mating partner before crossing, i.e., in progeny of *blr1*Δ (*hph* and *hph*-ΔN), but not in progeny of *ku70*Δ or *env1Δ* that comprise only one copy of full-length *hph* in their genome (Figure 6C and Additional File 1: Table A14).

To determine whether RIP could operate between two genetically unlinked *hph* alleles, we generated a *ku70*Δ*env1*Δ double mutant that had one deletion on ChII (env1Δ) and the other on ChIII (*ku70*Δ). Our data revealed that sexual crossing (n = 10) between *ku70*Δ*env1*Δ and a wild-type mating partner also resulted in C-to-T point mutations in progeny of these crosses (Figure 6C and Additional File 5).

In *Neurospora crassa*, sequences mutated by RIP showed skewed dinucleotide frequencies because of the sequence preference of RIP (CpA > CpT > CpG > CpC) [64]. By comparing the nucleotide sequences of the *hph* alleles in all F1 progeny (Appendix A4), we found that *Trichoderma reesei* displayed a different sequence preference of RIP, i.e., CpA ≈ CpG >> CpT > CpC (Additional File 1: Table A14).

We conclude that *Trichoderma reesei*, like *Neurospora crassa*, exhibits high homology pairing and RIP activities at a premeiotic stage before premeiotic DNA synthesis. Our results are thus of tremendous importance for industrial strain improvement.

### Repetitive features

The completeness of the QM6a and other 11 high-quality fungal genomes also allowed us to accurately survey genome-wide repetitive features and their correlation to RIP using the RepeatMasker search program (http://www.repeatmasker.org/). We were able to identify almost all Ty elements along the 16 chromosomes of *Saccharomyces cerevisiae* [65, 66] (Additional File 1: Table A15). Our results also confirm that the genome of *Neurospora crassa* accumulates fragmented and rearranged transposon relics, in particular *gypsy*-LTRs (long terminal repeats) and Tad-LINEs (long interspersed repeat elements) [30]. The fungal wheat pathogen *Mycophaerella graminicola* has a high copy number of *gypsy*-LTRs, copia-LTRs and Tad-LINEs [55]. *gypsy*-LTRs are highly overrepresented in the genomes of *Schizosaccharomyces pombe* [67], *Cryptococcus neoformans* [68] and *Coprinopsis cinerea* [69]. The *Candida glabrata* CBS138 genome contains very few (∼0.4%) repetitive sequences (http://www.candidagenome.org/)(Additional File 1: Table A16).

Intriguingly, the QM6a genome has the fewest transposon sequences among the six Group I fungal genomes (Additional File 1: Table A16). The majority of *copia*-LTRs (6/8), *gypsy*-LTRs (9/10), *CMC-EnSpm* (6/6) and *MULE-MuDR* (13/21) are located in longer AT-rich blocks. In contrast, most LINEs (16/18) are located in non-AT-rich regions (Additional File 1: Table A17). Neither centromeres nor telomeres show a preponderance of any particular type of transposable element (Additional File 1: Table A17). According to their RIP indices (Additional File 1: A13) and smaller size (Additional File 1: Tables A17 and A18), we conclude that almost all transposon sequences in QM6a are fragmented or rearranged transposon relics.

The copy numbers of transposon sequences in QM6a we report here (Additional File 1: Table A16) are much lower than those reported by Kubicek et al. using the QM6a-v2.0 draft [13]. Both we and Kubicek *et al*. used the RepeatMasker search program to search for repetitive sequences. It is important to point out that the limitation of this widely-used program is that it often generates many false-positive results. This is why we first applied it to the *Saccharomyces cerevisiae* genome to determine the optimum parameters for filtering the preliminary RepeatMasker data (see Materials and Methods). The number and locations of five different Ty elements in all of that 16 yeast’s chromosomes had been determined before [65, 66].

### Partitioning of gene clusters by AT-rich blocks

A hallmark of the QM6a-v2.0 draft genome is that a third of the 228 CAZyme genes are non-randomly distributed and form several CAZyme gene clusters. Several of the regions of high CAZyme gene density also contain genes encoding proteins involved in secondary metabolism. Accordingly, it has been proposed that gene clustering and/or coexpression might be evolutionarily advantageous for *Trichoderma reesei* in its competitive soil habitat or other natural environments [11, 70]. Using the QM6a-HiC draft genome as a reference, 20 CAZyme gene clusters and 42 SSCP gene clusters were identified in the QM6a-HiC genome draft [18]. All these gene clusters consisted of only 3-6 CAZyme and/or SSCP genes but, surprisingly, gene clustering did not influence gene expression.

To gain a better insight, we reexamined all these 62 CAZyme and SSCP gene clusters in QM6a-HiC [18]. Due to sequencing and assembly errors, some gene clusters are overlapped or even duplicated. The original 62 gene clusters locate to 46 chromosomal blocks in the complete QM6a genome. The majority of these gene clusters also contain new QM6a genes we annotate in this study (Additional File 2: Table B4). One gene cluster even contains a counterfeit gene (QM6a-v2.0 gene number 71245) [18].

Intriguingly, except for 5 short gene clusters (≤ 4 contiguous genes), all the other 41 gene clusters in the complete QM6a genome are divided into smaller compartments by AT-rich blocks (Additional File 2: Table B4). The first example of this phenomenon is a nitrate assimilation gene cluster and an annotated CAZyme gene cluster [18] immediately adjacent to *tel2R*. These two gene clusters were divided by *tel2R* and five interspersed AT-rich blocks into five smaller compartments (Figure 7A). The first (or rightmost) compartment consists of one hypothetical protein gene (TrB1976W) and three nitrate reductase genes (*nit3*, TrB1975W; *nit2*, TrB1974C; *nit6*, TrB1973C); transcripts of these genes were barely detectable (TPMs < 0.5) under glucose for 48 hours, then in straw for 24 hours and finally with the addition of glucose for 5 hours. The second compartment comprises a sole nitrate transporter gene (*nit10*, TrB1972C); *nit10* was slightly induced by straw and then repressed by the addition of glucose. The third compartment contains a β-mannosidase gene, a member of glycoside hydrolase family 2 (GH2, TrB1971C) and a hypothetical gene (TrB1970W). Compared to *nit10*, this β-mannosidase gene exhibited ∼∼5-fold greater induction by straw. The fourth compartment has only a β-1,4-glucuronan lyase gene (*trgL*, TrB1969WW). Like the nitrate reductase genes, *trgL* did not respond to glucose or straw substrate. Finally, the last (or leftmost) compartment harbors two CAZyme genes and a hypothetical protein gene (TrB1966). The two CAZyme genes are the glucuronoyl esterase *cip2* (TrB2050W) and a GH30 endo-β-1,4-xylanase (TrB2049C) (Figure 7A). These two genes were highly induced by straw and then repressed by the addition of glucose (Additional File 1: Table A19A).

**Figure 7.**
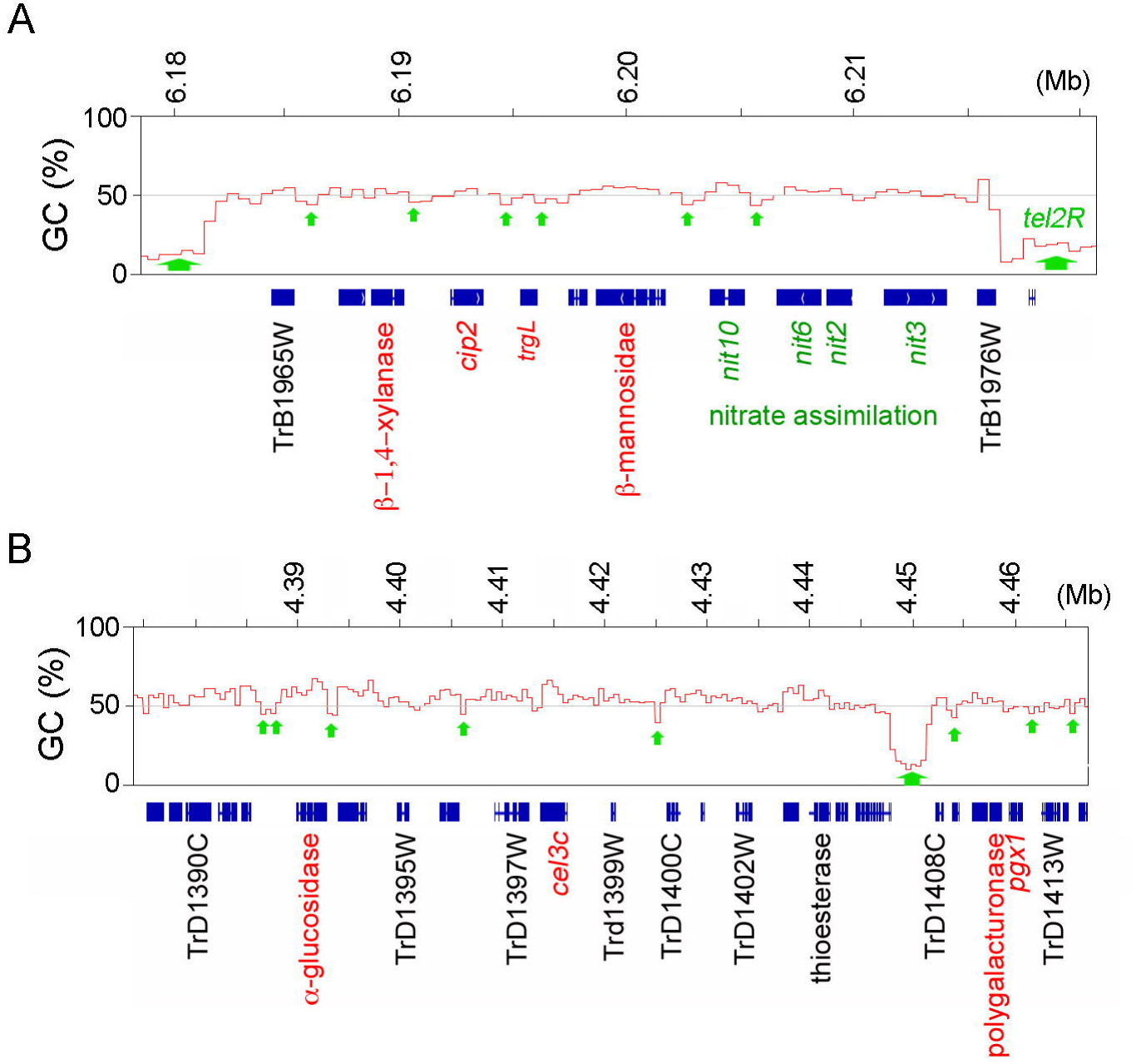
Partitioning of gene clusters by AT-rich blocks. The tracks represent the graphs of GC contents (window size 500 bp) of a gene cluster next to *tel2R* (A) and pericentromeric regions (B). All exons of all the predicted genes are indicated with blue squares, short AT-rich blocks by smaller green arrows, *tel2R* and long AT-rich blocks by larger green arrows, the CAZyme genes in red, the nitrate assimilation genes in dark green and the systematic names of representative QM6a genes in black.

The second example is a CAZyme gene cluster on ChIV with 22 genes. It is divided by eight AT-rich blocks into seven smaller compartments (C1-C7). The four CAZyme genes are located in different compartments; a GH31 α-glucosidase (TrD1393W) in C1, *cel3c* (TrD1398C) in C3, and a GH28 polygalacturonase (TrD1411C) and *pgx1* (TrD1412C) in C6 (Figure 7B). These four CAZyme genes were differentially regulated by straw and the addition of glucose (Additional File 1: Table A19B).

The third example is the CAZyme gene cluster close to *tel7R*. Six AT-rich blocks partitioned this gene cluster into four smaller compartments (C1-C4). The four CAZyme genes were allocated to three different compartments; the *egl2/cel5* endo-β-1,4-glucanase (TrG1193W) in C1, GH79 ß-glucuronidase (TrG1202W) and rhamnogalacturonyl hydrolase (TrG1204W) in C2, and the *cbh* cellobiohydrolase (TrG1206C) in C4. Only *cbh* was highly induced by straw in QM6a (Additional File 1: Table A19C).

Together, these results might explain why it was previously reported that gene clustering did not influence gene expression [18]. We propose that these smaller compartments are structurally and functionally similar to intrachromosomal loops or TADs.

It should be noted that occurrence of intergenic AT-rich blocks did not always result in differential gene expression. For example, a gene cluster with three contiguous CAZyme genes—the acetyl xylan estrease *gene axe1* (TrE0669C; acetyl xylan estrease), the cellulose-induced protein *cip1* (TrE0670C) and the β-1,6-Ν-acetylglucosaminyl transferase gene *egl4* (TrE0671C)—resides in the middle of ChV. These three CAZyme genes are separated by four AT-rich blocks into three smaller compartments, but they were all highly induced by straw and then repressed by the addition of glucose (Additional File 1: Table A19D). In this case, each gene becomes a partitioned functional unit (see Discussion). Simultaneous expression of the *axe1-cip1-egl4* triad might be independently controlled by other determinants, e.g., common transcription factors and/or similar chromosomal conformation.

## Discussion

*Trichoderma reesei* QM6a and its derivatives have been widely used for nearly four decades to produce plant cell wall-degrading enzymes and heterologous recombinant proteins. In this study, we have obtained a high-quality complete genome sequence of QM6a. We readily uncovered many sequencing, assembly and gap-closing errors in earlier draft genomes. The seven telomere-to-telomere QM6a chromosomes can be used as better scaffolds for comparative genomic analyses, not only with industrial strains but also other *Trichoderma reesei* wild isolates (e.g., CBS999.97) and other species in the same genus (e.g., *Trichoderma atroviride* and *Trichoderma virens*). Our results also revealed much new genomic information never provided by earlier draft genomes, including 7 centromeres, 14 telomeres, 2328 AT-rich blocks, 1630 newly-predicted genes, 37 centromere-encoded genes and 24 centromeric repeats. Therefore, our complete QM6a genome sequence provides a comprehensive roadmap for further studies of this economically-important fungus, including industrial strain improvements and elucidation of the functional relationships between sequences, gene products and genome organization.

The central finding of this study is that *Trichoderma reesei* has evolved a robust RIP system. Firstly, the QM6a genome contains the lowest overall copy number of transposons among six studied filamentous ascomycetes (including *Neurospora crassa*). Secondly, as in *Neurospora crassa*, sexual crossing readily induced C-to-T point mutations on both tandem and unlinked duplicated sequences in *Trichoderma reesei*. Thirdly, almost all 2349 AT-rich blocks in QM6a were predicted by the RIPCAL software program to be affected by RIP. Considerable evidence suggests that AT-rich blocks establish a link between DNA sequence and chromatin architecture. In *Neurospora crassa*, AT-rich blocks form constitutive heterochromatins and mediate intra- and inter-chromosomal contacts [46, 47]. In *Saccharomyces cerevisiae*, AT-rich blocks constitute the chromosome axes or the bases of chromosomal loops or TADs [35, 38-40]. In mammalian interphase chromosomes, the spatial distribution of AT-rich blocks (e.g., lamina-associated domains, LADs) and GC-rich blocks (e.g., TADs or chromosomal loops) are evolutionarily conserved. LADs preferentially interact with other LADs, whereas TADs exhibit more localized chromosomal domains [71]. Intriguingly, our results reveal that the rDNA locus of QM6a is surrounded by two interspersed AT-rich blocks. It would be of interest to further investigate whether and how these two interspersed AT-rich blocks are involved in preventing rDNA from being affected by RIP as well as in regulating nucleolar organization, rRNA transcription and rDNA copy homeostasis.

From the results of this study, we postulate that RIP does not function solely as a genome defense mechanism to diminish the potentially deleterious effects caused by the spread of transposable elements. It may also have important roles in reshaping the *Trichoderma reesei* genome. We demonstrate that the widespread interspersed AT-rich blocks lead to genome-wide partitioning of the gene clusters in QM6a. Our findings can readily account for why gene clustering does not affect gene expression in *Trichoderma reesei*. Mechanistically, RIP-mediated C-to-T mutations presumably can transform duplicated sequences in a CAZyme gene cluster into interspersed AT-rich blocks, thus dividing an ancestral gene cluster with multiple CAZyme or SSCP genes into multiple smaller compartments or TADs. Intriguingly, it has been reported previously that many pathogenic fungi (*Leptosphaeria maculans, Magnaporthe oryzae, Fusarium* spp.) comprise AT-rich blocks with RIP affected effector genes and transposable elements [72, 73] and that RIP is a potential factor in *Leptosphaeria maculans* in creating the rapid sequence diversification (i.e., of the effector genes) needed for selection pressure [74] and concerted epigenetic regulation of their expression [75]. Partitioning of gene clusters by AT-rich blocks may also help to control simultaneous expression of the rDNA locus (Figure 3) and of the three partitioned functional units in the *axe1-cip1-egl4* triad (Additional file 1: Table A19D). Further research will reveal how RIP provides evolutionary advantages to *Trichoderma reesei* and other filamentous ascomycetes (Pezizomycotina) to survive in natural environments and pathogenic conditions.

## Conclusion

The earlier genome drafts of QM6a and other *T. reesei* industrial strains have been useful in identifying key genes involved in several important biological processes. However, due to numerous sequencing and assembly errors, they are not suitable for several other studies, e.g., genome-wide determination of gene order, chromosome architecture and expression dynamics as well as chromosome engineering for genetic modification(s). The complete QM6a genome sequence provides an unprecedented opportunity to overcome those obstacles associated with earlier draft genomes. To avoid the limitations of working with incomplete datasets and the false leads that can come from trying to work with imperfect data, more caution should be exercised in utilizing the draft genome sequences solely determined by SGS and/or HiC for functional and comparative analyses.

## Materials and Methods

### Fungal growth, DNA preparation and pulsed-field gel electrophoresis (PFGE)

QM6a was inoculated on petri-plates with malt extract agar (MEA) medium at 25 °C until full asexual sporulation was observed (∼5 days). 2×10^8^ conidial spores were collected, and then inoculated in 50 mL potato dextrose medium (PDB) at 30 °C for 6 h. The germinate hyphae were harvested by centrifugation at 3000*g* for 5 min at room temperature and incubated in 2 mL lysing enzyme buffer [0.1 M KH_2_PO_4_ (pH 5), 1.2M Sorbitol, 5% lysing enzyme (Sigma, USA)] at 30 °C for 1.5 h. The protoplasts were harvested by centrifugation at 600*g* for 10 min at 4 °C, dissolved in 1.2 ml GUHCl solution (43% guanidine-HCl, 0.1M EDTA pH8.0, 0.15M NaCl, 0.05% Sarkosyl) at 65 °C for 20 min and then mixed with 6.4 mL ice-cold ethanol to precipitate the genomic DNA. The pellet was dissolved in 10X TE with 0.6 mg/ml RNaseH at 37 °C for 1hr and then in 0.4 mg/mL proteinase K at 65 °C for 1hr. The genomic DNA was purified with the phenol:chloroform:isoamyl alcohol (25:24:1) method and then recovered by standard precipitation with ethanol. Next, the quality of high molecular weight genomic DNA for Illumina MiSeq and PacBio sequencing was validated by PFGE. The genomic DNA was separated in a 1% agarose gel in 0.5x TBE buffer, using a CHEF DR II (Biorad) with 0.5x TBE running buffer, continuously refrigerated at 14 °C and 6 V/cm (current 110-125 mA) for 18 h. The Lambda DNA-MonoCut Mix (New England Biolabs, N3019S) was used as size marker. Visualization was performed after staining with ethidium bromide after the electrophoresis.

### DNA sequencing and de novo assembly

Illumina MiSeq sequencing was carried out at the DNA sequencing facility in the Institute of Molecular Biology, Academia Sinica (Taipei, Taiwan) in May 2016 (sequencing was coordinated by the author SYT). A shotgun paired-end library (average size of 550 bp) was prepared using the Illumina TruSeq DNA nano Sample Prep Kit. The Illumina library (including PCR amplification and quantification) was prepared automatically by the NeoPrep Library Prep System, and then sequenced on the Miseq platform (300 cycles, paired-end sequencing) with the Miseq Control software version 2.5.1 and Sequencing Analysis Viewer version 1.8.20. Sequencing data were sent to Illumina BaseSpace automatic analysis during running. For report version 2.2.9 with MiSeq 600 cycler V3 chemistry, 96.16% of clusters passed the filter and 78.32% of bases qualified higher than Q30 at 2×300 n.

For Pacbio continuous long read sequencing, high-quality genomic DNA was submitted to the Ramaciotti Centre for Genomics (University of New South Wales, Sydney, Australia). Sequencing was coordinated by Carolina Correa and Tonia Russel. High molecular weight DNA was sheared with g-TUBE (Covaris PN 520079), aiming at DNA fragments of about 20 kb. The library was constructed with a 20kb size-selected protocol using DNA Template Prep Kit 2.0 (PN 001-540-726), purified and further selected for long insert size with a 0.35X AMPure (AMPure PB PN 100-265-900) bead size selection. The library was sequenced on a PacBio RSII device using the reagents DNA/Polymerase Binding Kit P4 (PN 100-236-500), DNA Sequencing Reagent 2.0 PN (100-216-400), SMRT®Cell V3 (PN 100-171-800) and Magbead (PN 100-133-600), loading at 200 pM on the plate. Data were collected with Stage Start and 180 minute movies. Seven SMRT®Cell cells were used to generate 263,312 reads and 3,397,762,180 bases.

The Pacbio data were assembled with the SMRT Analysis Software v2.3.0 (http://www.pacb.com/products-and-services/analytical-software/smrt-analysis/). The assembly was performed with HGAP (3.0 protocol) with the following parameters: (1) PreAssembler Filter v1 (minimum sub-read length = 500bp, minimum polymerase read quality = 0.80, minimum polymerase read length = 100bp); (2) PreAssembler v2 (minimum seed length = 10,000bp, number of seed read chunks = 6, alignment candidates per chunk = 10, total alignment candidates = 24, min coverage for correction = 6); (3) AssembleUnitig v1 (Genome Size 34000000bp, target genome coverage = 30, overlap error rate = 0.06, minimum overlap = 100bp and overlap k-mer = 22); (4) BLASR v1 mapping of reads for genome polishing with Quiver (max divergence percentage = 30, minimum anchor size = 12). The final assembly contained nine unitigs (seven telomere-to-telomere chromosomes, a circular mitochondrial genome and a 1870bp short sequence) with a total 34,993,035 bp and approximately 87x genome coverage. N50 was 5,311,312 bp and max was 6,835,650 bp. The per-base Quality Value (QV) is higher than 45 and a little less than 50 (99.999% accuracy; average 48.8). A BLASTN search revealed that the circular mitochondrial genome (42,130 bp) was completely identical in nucleotide sequence with the QM9414 mitochondrial genome (accession: NC_003388.1) [21]. The shortest unitig (1870 bp) is likely contaminating DNA because no Illumina MiSeq reads were mapped to its sequence.

### Mapping of Illumina reads over three different genomic drafts

The two runs of Illumina MiSeq with 300-bp paired-end reads were combined into one. The forward and reverse data were 6.8 Gb. Reads were preprocessed with Trimmomatic V0.32 (http://www.usadellab.org/cms/?page=trimmomatic) to trim and remove reads (33.15%) that fell below a quality score threshold of 30 (Q30) and were shorter than 30bp. Next, CLC Genomics Workbench 7.5 (http://www.clcbio.com/blog/clc-genomics-workbench-7-5/) was used to map the qualified reads to three different QM6a genomic drafts: (1) QM6a-v2.0 (http://genome.igi.doe.gov/Trire2/Trire2.home.html) plus the complete mitochondrial genome [20]; (2) the HiC draft genome [16] plus the complete mitochondrial genome [20]; (3) the PacBio *de novo* assembly (this study). The parameters used for mapping were (1) Mismatch cost = 3; (2) Insertion cost = 2; (3) Deletion cost = 2; (4) Length fraction = 0.8; (5) Similarity fraction = 0.8. The results are listed in Table S4. There were some differences between the PacBio and Illumina platforms, so the mapping results were used to extract a consensus sequence to adjust the bases to a final version of the chromosome sequences. We defined a threshold=2 to identify low coverage regions. For low coverage regions (i.e. threshold ≥2), sequences from the Illumina Miseq platform were used to construct the consensus sequence.

The seven telomere-to-telomere chromosomes were categorized with Roman numerals (ChI-ChVII), from the largest to the smallest. Artemis was used to determine the percentage of G+C in 500 bp non-overlapping windows. The centromere of each chromosome was their longest interspersed AT-rich block. The EMBOSS revseq software tool (v6.5.7; http://www.bioinformatics.nl/cgi-bin/emboss/revseq) was used to reverse and complement the nucleotide sequences of ChII, ChIV and ChVI, respectively, so that all seven chromosomes had a shorter (left) arm at 5′ termini and a longer (right) arm at 3′ termini. The final sequences were first compared to the original version using BLAST searches (Additional File 1: Table A1) before being submitted to NCBI (accession numbers CP016232-CP016238).

### Genome reannotation

The MAKER2 v2.31.8 (http://www.yandell-lab.org/software/maker.html) genome annotation pipeline [74] was applied for genome reannotation. Firstly, all (28748) *T. reesei* protein sequences from NCBI were used for *ab initio* gene predictions. Secondly, the Augustus v3.0.3 gene prediction software (http://augustus.gobics.de/) [75] was also used to predict new genes. All proteins from *Neurospora spp*. and *Fusarium spp*. were used for Augustus training to impose constraints on the predicted gene structure, including splice sites, translation initiation sites and stop codons. Thirdly, we isolated QM6a poly(A) RNA and performed a MiSeq RNA sequencing experiment. The resulting mRNA reads (trimming quality=30, min length=20bp) were applied to Trinity (v2.0.6; https://github.com/trinitvrnaseq/trinitvrnaseq/wiki) for *de novo* transcriptome assembly. The expressed sequence tag (EST) results predicted by “Maker” were used to identify new genes. Finally, we integrated all the ESTs and protein sequences from the two publicly-available databases (QM6a-v2.0 and RUT-C30-v1.0; at JGI) as well as all the ESTs assembled from two published transcriptome datasets [24, 25].

#### A. QM6a-v2.0

(http://genome.jgi.doe.gov/Trire2/Trire2.home.html)
TreeseiV2_FilteredModelsv2.0.transcripts.fasta
TreeseiV2_FilteredModelsv2.0.proteins.fasta

#### B. RUT-C30-v1.0

(http://genome.jgi.doe.gov/TrireRUTC30_1/TrireRUTC30_1.home.html)
TrireRUTC30_1_GeneCatalog_transcripts_20110526.nt.fasta
TrireRUTC30_1_GeneCatalog_proteins_20110526.aa.fasta.

#### C. The Illumina HiSeq 2000 sequencing reads from QM9414 treated with cellulose (24, 48 and 72 hours), sophorose (2, 4 and 6 hours) and glucose (24 and 48 hours) [25]

GSE53629 (http://www.ncbi.nlm.nih.gov/geo/query/acc.cgi?acc=GSE53629)
SRR1057947: QM9414 Cellulose replication 1
SRR1057948: QM9414 Cellulose replication 2
SRR1057949: QM9414 Cellulose replication 3
SRR1057950: QM9414 Sophorose replication 1
SRR1057951: QM9414 Sophorose replication 2
SRR1057952: QM9414 Sophorose replication 3
SRR1057953: QM9414 Glucose replication 1
SRR1057954: QM9414 Glucose replication 2
SRR1057955: QM9414 Glucose replication 3

The Illumina-HiSeq .sra files were converted into fastq using the fastq-dump program of the SRA Toolkit. Trimmomatic v0.32 [76] was used to trim and remove reads that fell below a quality score threshold of 30 (Q30) and that were shorter than 10bp. The Cufflinks pipeline tools (http://cole-trapnell-lab.github.io/cufflinks/) were downloaded for transcriptome assembly and differential expression analysis, including bowtie2-build (bowtie v2.2.3), tophat2 (v2.0.13) and cufflinks (v2.2.1). For read alignment, the parameters of tophat2 were: (1) min-intron-length: 20 bp; (2) max-intron-length: 5000bp (to reflect introns and splicing elements of five diverse fungi); and (3) No transcript GTF (Gene Transfer Format) file was provided to guide assembly. There were ∼95% trimmed paired reads that could be aligned to the QM6a complete genome. After each QM9414 RNA-seq raw dataset was assembled into transcripts, the cuffmerge program was used to merge all GTF files into one. The gffread program was used to extract transcript sequences.

#### D. The SOLiD sequencing reads from QM9a grown first in glucose for 48 hours, then in straw for 24 hours and finally with the addition of glucose for 5 hours [24]

GSE44648 (http://www.ncbi.nlm.nih.gov/geo/query/acc.cgi?acc=GSE44648)
SRR764963: QM6a Glucose 48 h replication 1
SRR764964: QM6a Glucose 48 h replication 2
SRR764965: QM6a Glucose 48 h replication 3
SRR764966: QM6a Straw 24 h replication 1
SRR764967: QM6a Straw 24 h replication 2
SRR764968: QM6a Straw 24 h replication 3
SRR764969: QM6a Straw + Glucose 5 h replication 1
SRR764970: QM6a Straw + Glucose 5 h replication 2

The SOLID .sra files were converted into csfasta and QV.qual using the abidump program of the SRA Toolkit. For transcriptome assembly and quantification, bowtie-build (bowtie v1.1.0), tophat2 and cufflinks were used. All parameters were as for the QM9414 dataset, except the colorspace option was used for the bowtie1 program. There were ∼33% reads that could be aligned to the QM6a complete genome.

Next, we performed gene filtering to finalize all predicted protein sequences, the filtering order was all *Trichoderma reesei* protein sequences from NCBI > *Neurospora spp*. and *Fusarium spp*. (Augustus v3.0.3) > *de novo* assembly QM6a-RNA (Trinity v2.0.6) > QM6a v2.0 + RutC-30 v1.0 + QM6a-RNA (SOLiD) + QM9414-RNA (Illumina HiSeq 2000).

We also manually integrated almost all the annotation results reported by the *Trichoderma* research community [4, 18, 24, 25]. We annotated 10786 predicted genes that covered almost all previously annotated genes in QM6a-v2.0 (9105 out of 9129) and RUT-C30-v1.0 (9717 out of 9852), respectively. Of these predicted QM6a genes, 904 were previously considered to be RUT-C30-specific. We successfully annotated several evolutionarily-conserved DNA repair genes, including *rad50, sae2/com1*/CtIP, *rad51, rad57, srs2* and *pif1* (Additional File 2: Tables B1 and B2). These genes were not annotated in QM6a-v2.0 (JGI), RUT-C30-v1.0 (JGI) and QM6a-HiC [18].

Finally, the BLASTP program was applied to compare our final predicted protein sequences to several publicly-available protein databases, including NCBI nonredundant (nr) database, Universal Protein Resource (Uniprot) Prot v2.0, *Neurospora crassa* database (Broad Institute), *Fusarium fujikuroi* (JGI), *Sordaria macrospora* (NCBI), *Saccharomyces cerevisiae* (SGD), and *Schizosaccharomyces pombe* (PomBase and NCBI). The e-values of all BLAST results were < 1.0 × 10^−5^. The Retrieve/ID mapping tool (Uniprot) was used to map gene ID (Uniprot sprot v2.0 and NCBI BLAST result) and to determine gene ontology (GO) (Additional File 2: Tables B2).

Next, the bowtie-tophat program was used to align all QM6a (SOLiD) and QM9414 (Illumina MiSeq) RNA reads with the following parameters: (1) min-intron-length: 20bp; (2) max-intron-length: 5000 bp (because it was reported previously that the ranges of intron length in *Saccharomyces cerevisiae, Aspergillus nidulans* and *Neurospora crassa* are 52-1002 bp, 27-1903 bp and 46-1740 bp, respectively [76]); and (3) a transcript GTF (Gene Transfer Format) file was provided. To calculate read counts for each transcript, we applied featureCounts [77]. The values of Transcripts Per kilobase Million (TPM) [26] were used as expression values: TPM = (individual gene RPK/the sum of all RPKs) x 10^6^, whereas RPK (Reads per kilobase) = (read counts/transcription length) (Additional File 2: Tables B1 and Additional File 3).

### Repeat Modeler and RepeatMasker

Novel repeat elements were identified by Repeat Modeler-1.0.4 (http://www.repeatmasker.org/RepeatModeler.html) with default parameters. RepeatMasker (version 4.0.6) and the Repbase Library (http://www.repeatmasker.org) were used to scan 12 different fungal genomes for interspersed repeats and low complexity DNA sequences. The output of the program is a detailed annotation of the repeats that are present in the query sequence, as well as a modified version of the query sequence in which all the annotated repeats have been masked (default: replaced by Ns). To obtain high-confidence data, we first analyzed the genome sequences of *Saccharomyces cerevisiae* because the number and location of five different Ty elements in all its 16 yeast chromosomes had been reported previously [65, 66]. When the preliminary RepeatMasker data were filtered with two parameters (length >= 140, Smith–Waterman local similarity scores >= 450), the final data (Additional File 1: Table A15) were quite consistent with the known results [65]. The same parameters were then applied to scan all other studied fungal genomes (Additional file 1: Table A16).

### RIP and RIPCAL

The latest RICAL program (https://sourceforge.net/projects/ripcal/) was downloaded [58, 59]. An initial analysis was done to predict if the complete QM6a genome sequence had been mutated by RIP. All strains used in this study had been previously described [10]. The methods for sexual crossing, single ascospore isolation and preparation of genomic DNA have also been described before [8, 10]. The full-length *hgh* cassettes in all F0 parental strains and representative F1 progeny were amplified by PCR and analyzed by Sanger sequencing technology. All the nucleotide sequences of primers and the *hgh* cassettes are listed in Additional file 1: Table A9 and Additional file 5, respectively. The RIPCAL dinucleotide frequency and EMBOSS compseq tool were used to determine the values of two RIP indices, TpA/ApT and [(CpA + TpG) / (ApC + GpT)], respectively.

## Declarations

### Authors’ contributions

WCL and CLC isolated the fungal genomic DNA for sequencing and conducted the sexual crossing experiments. CHH, WCL and YCC carried out bioinformatic analyses. SYT performed Illumina Miseq sequencing experiments. TFW conceived and designed the experiments, and analyzed the data. TFW and WCL wrote the paper. All authors read and approved the final manuscript.

## Acknowledgments

We thank Paul Hsu at the IMB Bioinformatics Service Core, Yu-Tang Huang at the IMB Computer Room and the IMB Genomics Core for technical support, and John O’Brien for English editing. TFW would like to thank Frédérique Bidard-Michelot (IFP Energies Nouvelles, Paris, France) for personal communications on unpublished results of RUT-C30 genome assembly and a common consensus for QM6a chromosome numbering, ordering and orientation.

## Availability of data and materials

- All data generated or analyzed during this study are included in this published article and its supplementary information files. Data deposition: BioProject (PRJNA325840), BioSample (SAMN05250858), QM6a genomic DNA illumina Mi-seq (SRR4417032), QM6a Poly(A) RNA illumina Mi-seq (SRR5229930) and the nucleotide sequences of seven complete QM6a chromosomes (CP016232-CP016238).

## Ethical approval

This study needed no ethical approval and had no experimental research on humans.

## Additional files

**Li et al. (TFW) Additional file 1.pfd Additional File 1: Table A1.** Preliminary assembly results obtained by the Hierarchical Genome Assembly Process (HGAP 3.0). **Table A2.** Error corrections of the seven PacBio unitigs using the Illimina-MiSeq reads. **Table A3.** Characteristics and assembly of the seven QM6a chromosomes. **Table A4.** Mapping of all trimmed paired-end Illumina MiSeq reads to three QM6a genome drafts. **Table A5.** The MiSeq reads mapped to the QM6a-v2.0 draft genome. **Table A6.** Sixteen false predicted genes in QM6a-v2.0. **Table A7.** The QM6a-v2.0 draft genome compared to the complete QM6a genome sequence. **Table A8.** The HiC draft genome compared to the complete QM6a genome sequence. **Table A9.** PCR primers. **Table A10.** The complete QM6a genome sequence versus the RUT-C30-v1.0 draft genome. **Table A11.** Gene Ontology of some newly-predicted genes. **Table A12.** Sequencing quality of 18 different fungal genomes. **Table A13.** RIP indices of various sequences in QM6a and *Neurospora crassa*. **Table A14.** Repeat-induced C-to-T mutations observed in the *hph* alleles in all F1 progeny. **Table A15**. *Saccharomyces cerevisiae* Ty elements by chromosome. **Table A16.** Transposable elements in 12 well-assembled fungal genomes. **Table A17.** Repetitive sequences in different chromosomal regions. **Table A18.** Repetitive sequences in seven QM6a chromosomes. **Table A19.** Partitioning of four gene clusters by the AT-rich islands. The TPM values in glucose (48 hrs), in straw (24 hrs) and in straw (24hrs) then glucose (5 hrs) are shown [24].

**Li et al. (TFW) Additional file 2.xlsx Additional File 2: Table B1:** Genome annotation of the complete QM6a genomes, including AT-rich blocks, predicted genes, repetitive features, gene ontology (GO), gene names in *Trichoderma reesei, Saccharomycetes cerevisiae and Neurospora crassa*, as well as gene identity numbers in QM6a-v2.0 and RUT-C30-v1.0. (**Columns A-H**) All the genes that were not annotated in QM6a-v.20 and Rut-C30-v1.0 are highlighted with yellow background, whereas those only annotated in Rut-C30-v1.0 are highlighted with yellow-green background. (**Columns O-AE**) Comparative transcriptomic analysis of all QM6a genes in different carbon sources using two published transcriptome datasets: (1) The SOLiD sequencing reads from QM9a grown first in glucose for 48 hours, then in straw for 24 hours and finally with the addition of glucose for 5 hours [24]; (2) The Illumina HiSeq 2000 sequencing reads from QM9414 treated with cellulose (24, 48 and 72 hours), sophorose (2, 4 and 6 hours) and glucose (24 and 48 hours) [25]. The TPM values are shown. **Table B2:** Annotation of the complete QM6a genome. All the annotated QM6a genes are compared to those in QM6a-v2.0, Rut-C30-v1.0, all *Trichoderma reesei* proteins from NCBI, the QM6a-HiC annotation results by Druzhinina et al. [18], the annotation results by Schmoll et al. [4], as well as the BLAST results from Uniprot, NCBI non-redundant (nr) database, *Fusarium fujikuroi* (JGI), *Sordaria macrospora* (NCBI), *Saccharomyces cerevisiae* (SGD) and *Schizosaccharomyces pombe* (PomBase and NCBI). The e-values of all BLASTP results were < 1.0 × 10^−5^. **Table B3:** TPM values of all new QM6a genes revealed by two publicly-available transcriptome datasets [24, 25]. **Table B4:** Locations of 62 previously identified CAZyme and SSCP gene clusters [18] in 42 chromosomal blocks of the complete QM6a genome. All previously annotated QM6a-v2.0 or QM6a-HiC genes in these gene clusters are indicated in blue. AT-rich islands in these 42 chromosomal blocks are indicated in red. All new QM6a genes we identified in the complete QM6a genome are indicated in darkgreen.

**Li et al. (TFW) Additional file 3.xlsx Additional File 3: Table C1.** Straw-induced QM6a-v2.0 genes identified by Ries et al. [24] and their TPM values. **Table C2.** Straw-induced QM6a-v2.0 genes not identified by Ries et al. [24] and their TPM values. **Table C3.** New QM6a genes that are highly induced (≥ 20-fold) by straw and their TPM values. **Table C4.** New QM6a genes that are significantly induced (5-20fold) by straw and their TPM values. **Table C4.** Two new QM6a genes that are highly induced by cellulose and their TPM values. **Table C5**. Nine new genes having higher TPM values in QM6a than in QM9414. **Table C6.** Centromere-encoded genes and their TPMs in QM9414.

**Li et al. (TFW) Additional file 4.pdf Additional File 4:** Sequence alignments of the 24 centromeric repeats

**Li et al. (TFW) Additional file 5.pdf Additional File 5:** Nucleotide sequences of the *hgh* alleles from all strains listed in Figure 7.

